# Proteomic Analysis of Rat Serum Revealed the Effects of Chronic Sleep Deprivation on Metabolic, Cardiovascular and Nervous System

**DOI:** 10.1101/340000

**Authors:** Bo Ma, Jincheng Chen, Yongying Mu, Bingjie Xue, Aimei Zhao, Daoping Wang, Dennis Chang, Yinghong Pan, Jianxun Liu

## Abstract

Sleep is an essential and fundamental physiological process that plays crucial roles in the balance of psychological and physical health. Sleep disorder may lead to adverse health outcomes. The effects of sleep deprivation were extensively studied, but its mechanism is still not fully understood. The present study aimed to identify the alterations of serum proteins associated with chronic sleep deprivation, and to seek for potential biomarkers of sleep disorder mediated diseases. A label-free quantitative proteomics technology was used to survey the global changes of serum proteins between normal rats and chronic sleep deprivation rats. A total of 309 proteins were detected in the serum samples and among them, 117 proteins showed more than 1.8-folds abundance alterations between the two groups. Functional enrichment and network analyses of the differential proteins revealed a close relationship between chronic sleep deprivation and several biological processes including energy metabolism, cardiovascular function and nervous function. And four proteins including pyruvate kinase M1, clusterin, kininogen1 and profilin-1were identified as potential biomarkers for chronic sleep deprivation. The four candidates were validated via parallel reaction monitoring (PRM) based targeted proteomics. In addition, protein expression alteration of the four proteins was confirmed in myocardium and brain of rat model. In summary, the comprehensive proteomic study revealed the biological impacts of chronic sleep deprivation and discovered several potential biomarkers. This study provides further insight into the pathological and molecular mechanisms underlying sleep disorders at protein level.

## Introduction

Sleep is an essential and fundamental physiological process that plays crucial roles in the balance of psychological and physical health in almost all animals^[1]^. Accumulating evidence has demonstrated that sleep and wakefulness are critical to the cellular homeostasis associated with energy metabolism, synaptic potentiation, and responses to cellular stress^[2]^. In modern time, an increasing number of people have insufficient sleep and suffer from chronic sleep deprivation (CSD)^[3, 4]^.

CSD is a medical condition representing loss of sleep and shortness of sleep duration^[5]^. Clinical and experimental studies suggest that sleep loss may increase the risk of cardiovascular disorders, obesity, oxidative stress, diabetes and metabolic syndrome^[1, 2, 6, 7]^. Several metabolic alterations such as increased energy expenditure and intensified catabolism, which may result in weight loss and other metabolic dysfunctions, have been shown in rat models of paradoxical sleep deprivation^[8–10]^. CSD is associated with several cardiovascular dysfunctions and sleep duration was suggested to be an independent predictor for morbidity and mortality of cardiovascular disease^[11]^. In addition, SD may affect the cardiac autonomic nervous system leading to hypertension^[12–15]^. Prolonged wakefulness may also lead to dysfunction of immune system^[16]^. The results from the controlled SD experiments demonstrated that SD may increase lymphocyte activation and induce production of proinflammatory cytokines^[17]^. The increased inflammation factors such as IL-1, IL-2, TNF-α, C-reactive protein, and the augmented activities of monocytes, neutrophils, phagocytic cells and NK cells are believed to be involved in the extent of infarct and severe apoptosis in ischemic injury^[18]^. Furthermore, it has been reported that SD could disrupt secretion of hormones (e.g., thyroid stimulating hormone) and affect neuroendocrine function in patients with coronary heart disease. SD could also disturb melatonin rhythm, renin angiotensin system and metabolite rhythms^[5, 16]^.

Current proteomics technologies based on liquid chromatography coupled with tandem mass spectrometry (LC-MS/MS) have been widely employed to identify and quantify unique proteins and peptides in complex biological matrices such as serum^[19, 20]^. Proteomic analyses of serum are highly informative in detecting changes in protein levels and to determine the correlations of these clinically relevant biomarkers with various diseases^[21–25]^. A global survey of the changes in protein abundance in response to CSD would provide unique and meaningful insight into its biological mechanisms. Several previous studies based on proteomic technique of difference in gel electrophoresis (DIGE) or LC-MS were conducted to monitor the changes in protein abundance in brain tissues including astrocytes, cerebral cortex, thalamic and basal forebrain after sleep deprivation^[26–30]^. The results of these studies suggested that various biological processes including cell signaling, cytoskeletal, energy metabolism, exocytosis, mRNA processing/trafficking, neuronal transmission, neuronal plasticity, gliotransmission and vesicle trafficking are associated with CSD. In particular, energetics/metabolism and neuronal transmission were consistently suggested to be linked to CSD based on the brain tissue or cell line proteomic analyses. However, changes of serum protein in CSD using a LC-MS proteomic approach have not been reported.

In the current study, we aimed to analyze the serum proteome to identify the changes in protein targets potentially associated with CSD. The serum proteins were quantified to identify the changes in protein abundance in rat models with and without CSD.

## Materials and methods

### Experimental Design

Figure 1 shows the workflow of the serum proteomic profiling and analysis. Briefly, all the rats were randomly divided into 2 subgroups (n=12): the normal (N) group and chronic sleep deprivation (CSD) group. After 6 weeks, the blood was collected and the serum was obtained. Four serum samples from the same subgroup were combined and prepared for LC-MS analysis following the experimental procedures stated below. Peptides of each sample were then analyzed using a label-free quantitative (LFQ) proteomic technology. Proteins with differential abundance (DAPs) were analyzed using bioinformatics soft and online database. Several candidate proteins were identified and verified using western blot and parallel reaction monitoring (PRM) analyses. Finally, the potential biomarkers were validated by LC-MS analysis using myocardial and brain tissue of rat models.

**Figure 1.**
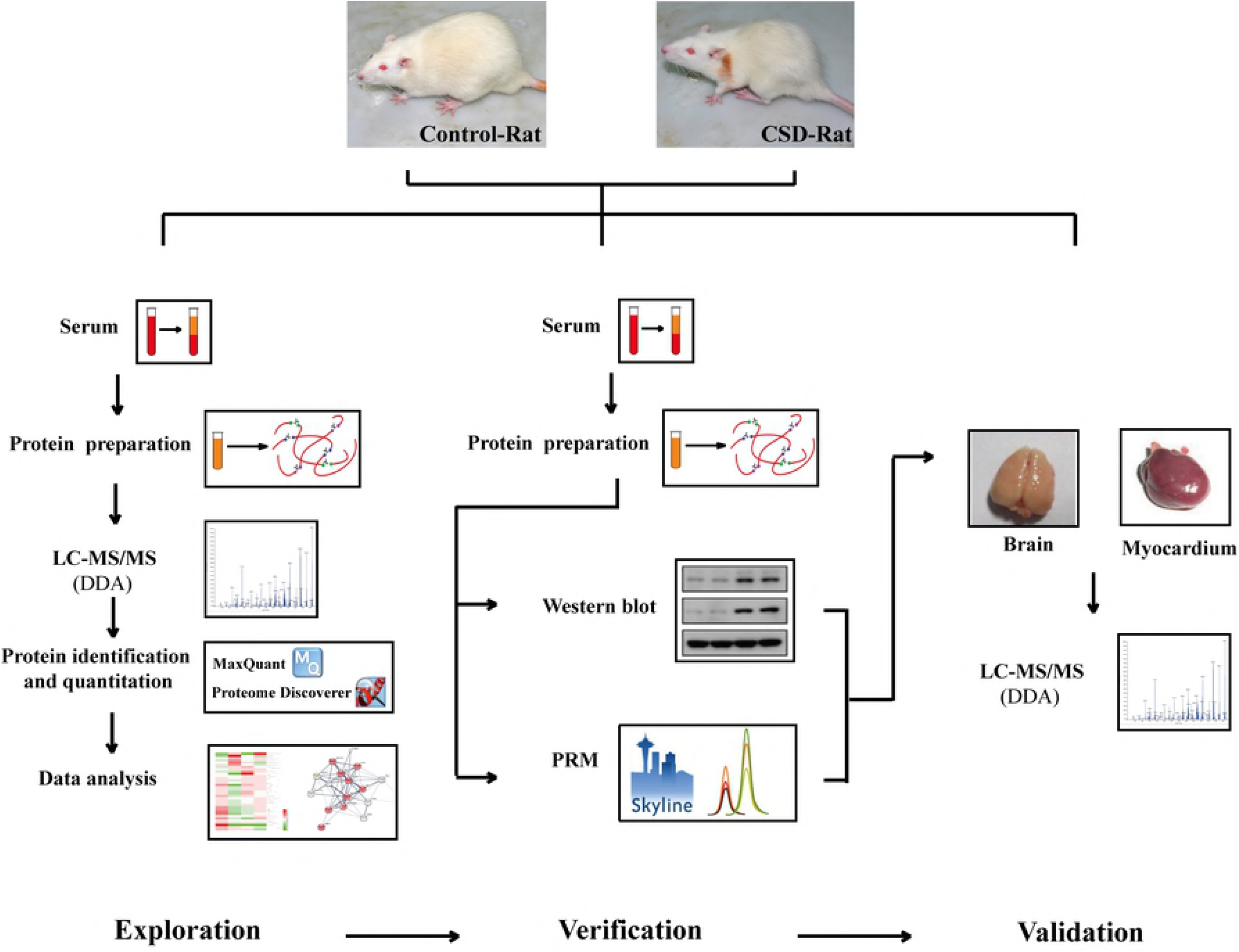
Workflow of the proteomic profiling and data analysis. Serum samples of normal (N, n=12) group and chronic sleep deprivation (CSD, n=12) group were analyzed by label-free quantitative proteomics methods. Proteins with differential abundance (DAPs) were analyzed using bioinformatics soft and online database. Several candidate proteins were identified and verified using western blot and LC-PRM analyses. Finally, the potential biomarkers were validated by LC-MS analysis using myocardial and brain tissue samples of rat models.

### Animal, grouping and chronic partial sleep deprivation

A total of 24 male Sprague-Dawley rats (Beijing Vital River Laboratory Animal Technology) with initial body weight of 180 ± 20 g, were randomly divided into two groups: control group (N group) and chronic sleep deprived group (CSD group). The animals in the CSD group were subject to sleep deprivation for 16 hours (16:00-8:00) per day over 6 weeks by the “flower pot” technique^[31–33]^. Briefly, animals were placed into regular container separately, and each CSD animal was placed on a round platform (diameter: 40-60 mm, height: 45-55 mm, surface was 5-10 mm above the water level). 6 platforms are situated in a rectangle container (600 mm × 450 mm) filled with room-temperature water, and the distance between two platforms is 150 mm. During sleep deprivation, muscle hypotonia caused animals to fall into the water, forcing them to climb back on the platform and remain awake. (The device has been under the patent substantive examination in China, NO: 201611090542.8.) Animals in the N group were placed in the same rectangle container without water to allow the animals to sleep under the same conditions. Animals were transferred to home cages for the remaining 8 h/day (8:00-16:00). All animals were housed in individual containers under standard conditions throughout the experiments and were maintained on a 12:12 light/dark cycle. The study was approved by the Ethics Committee of Xi Yuan Hospital, China Academy of Chinese Medical Sciences and was conducted in accordance with the ethical principles of animal use and care.

### Serum and tissue samples preparation and digestion

After 6 weeks of CSD, the animals in both N and CSD groups were sacrificed and the blood, myocardium and cerebral tissue were collected. The blood was immediately centrifuged at 1,500 g for 30 min to obtain serum. The serum samples were stored at – 80 °C until the assays were performed. Albumin and IgG of the serum was depleted using a ProteoExtract Albumin/IgG Removal Kits (Merck, Cat. No. 122642). For myocardial and cerebral tissue, each sample (100mg) was placed in a sample tube containing of 1ml lysis buffer (NaCl 150 mmol/L, 25 mM Tris-Cl pH 7.4, 1% TritonX-100, 0.1% SDS, 1 mM EDTA, 1% Nonidet P-40, 50 μg/ml PMSF). The sample tube was incubation for 1 h at 4 °C and lysed by ultrasound for 9 cycles of 20 seconds on ice. Then the tissue debris was removed by centrifugation at 15,000 rpm for 20 min at 4 °C, and the supernatant was collected.

Equal volume of the serum or tissue sample was treated with 5 mM of Tris (2-carboxyethyl) phosphine (TCEP, Sigma, Cat. No. C4706) for 30 min at 37 °C followed by 10mM of iodacetamide (IAM, Sigma, Cat. No. I1149) for 30 min at 37 °C. The proteins were then digested with trypsin (Promega, Cat. No. V5111, enzyme/protein ratio of 1:25 w/w) at 37 °C for rocking overnight. Undigested proteins and trypsin were removed by filtering the mixture through filters, the cutoff value of which was 10 kDa or more (Millipore).

### LC-MS/MS analysis

The peptides for DDA and PRM analysis were analyzed by nanoflow liquid chromatography-tandem mass spectrometry using a Q-Exactive plus mass spectrometer (Thermo, USA). The eluted peptides were separated on a pre-column packed with C18 Luna beads (3 μm diameter, 100 Å pore size; Thermo Fisher Scientific 164946) coupled to an analytical column packed with C18 Luna beads (3 μm diameter, 100 Å pore size; Thermo Fisher Scientific 164568). The binary solvent system was made up of 99.9% water and 0.1% formic acid (solvent A), and 99% acetonitrile and 0.1% formic acid (solvent B). Subsequently, the peptides were eluted with a linear gradient from 4% B to 35% B in 90 min with a constant flow rate of 400 nl/min. Eluted peptides were analyzed by DDA and PRM methods. Each experimental sample was analyzed in triplicate. A “wash column method” was executed between samples to avoid protein retaining in the column.

### Protein identification and quantitation

The resulting DDA Data were used for proteins identification and quantitation using Proteome Discoverer software (version 2.1, Thermo Scientific) and MaxQuant software (version 1.5.6.5)^[34]^. The mass spectrometry raw data was searched against the UniProt Rattus norvegicus protein database (downloaded February 2017). For the searches, precursor mass tolerance of 20 ppm, fragment mass tolerance of 0.1 Da and two missed cleavage were allowed. Variable modifications were set to: oxidation (M) and acetylation (protein N-term). Fixed modifications were set for carbamidomethylation of cysteine and variable modifications were set for oxidation of methionine. Protein level 1% FDR was set to filter the result. Details of the LC-MS data of rat serum, cerebral and myocardium tissues can be found in supplementary data 1-3.

Skyline software was used for PRM analysis. A scheduled (3-min window) inclusion list consisted of m/z of precursor peptides of interest and corresponding retention times was generated and used for scheduled PRM analyses. As described below, the final PRM analyses monitored the peptides from the 4 biomarker proteins. Peak area of the most intense product ions was considered for quantification. The product ion signals that showing interferences and did not match the retention time of the other monitored ions were excluded. The list of peptides targeted by PRM acquisition can be found in supplementary Table s2.

### Bioinformatics analysis

The information of the biological processes and molecular functions of the proteins was obtained from Gene Ontology (http://www.geneontology.org/) and UniProt Database. Heat map of the proteins with differential abundance was generated using Heml heat map illustrator program (version 1.0.3). Protein–protein interaction networks were analyzed by the STRING (Search Tool for the Retrieval of Interacting Genes/Proteins) system 10.5 (http://stringdb.org/).

### Echocardiography evaluation

Rat transthoracic echocardiograph was detected using VisualSonics Vevo2100 (VisualSonics, Canada) on the 6th week. Briefly, rats were anesthetized by 2-3% isoflurane and the anesthesia was maintained by 2% isoflurane with a nosecone on a heated platform. M-mode tracings of left ventricular short-axis views were recorded when the heart rate maintained at 330-380 bpm. Left ventricular ejection fraction (LVEF), fraction shortening (FS) and cardiac output (CO) were measured.

### Western blot

Protein expression levels were determined by Western blotting analysis. Equal amount of serum proteins (50μg) were separated on a SDS-PAGE gel and transferred to a polyvinylidene fluoride membrane. Membrane was blocked with TBST (20 mM tris-HCl, pH 7.6, 150 mM NaCl, and 0.1%tween-20), containing 5% BSA for 1 h, and then incubated with rabbit polyclonal primary antibody (Proteintech, 1:1000) at 4°C overnight, followed by incubating in horseradish peroxidase (HRP)-conjugated goat anti-rabbit secondary antibodies (Jackson, Cat. No. 111-035-003, 1:5000). Signals were detected using enhanced chemiluminescence (ECL) substrate (Proteintech, Cat. No. B500012), and imaged using the Tanon-5200 Chemiluminescent Imaging System (Tanon Science & Technology). As the shortage of GAPDH and β-actin in serum, coomassie blue staining of proteins was used to quality control of membrane transfer. Data are reported as representative results from at least three independent experiments.

### Statistical analysis

Data were shown as mean ± standard error for normally distributed values. All calculations were performed with GraphPad Prism program. A *P* value of less than 0.05 was considered statistically significant.

## Results

### Summary statistics of LC-MS/MS analysis

From the serum samples of N-and CSD-groups, LC-MS/MS analyses revealed a total of 309 non-redundant proteins with 297 and 298 proteins in N and CSD-groups, respectively. Of these proteins, 286 were found in both groups, constituting 92.6% of the total proteins identified (Figure 2A). 11 (3.6%) and 12 (3.9%) proteins were only found in the N and CSD-groups respectively.

**Figure 2.**
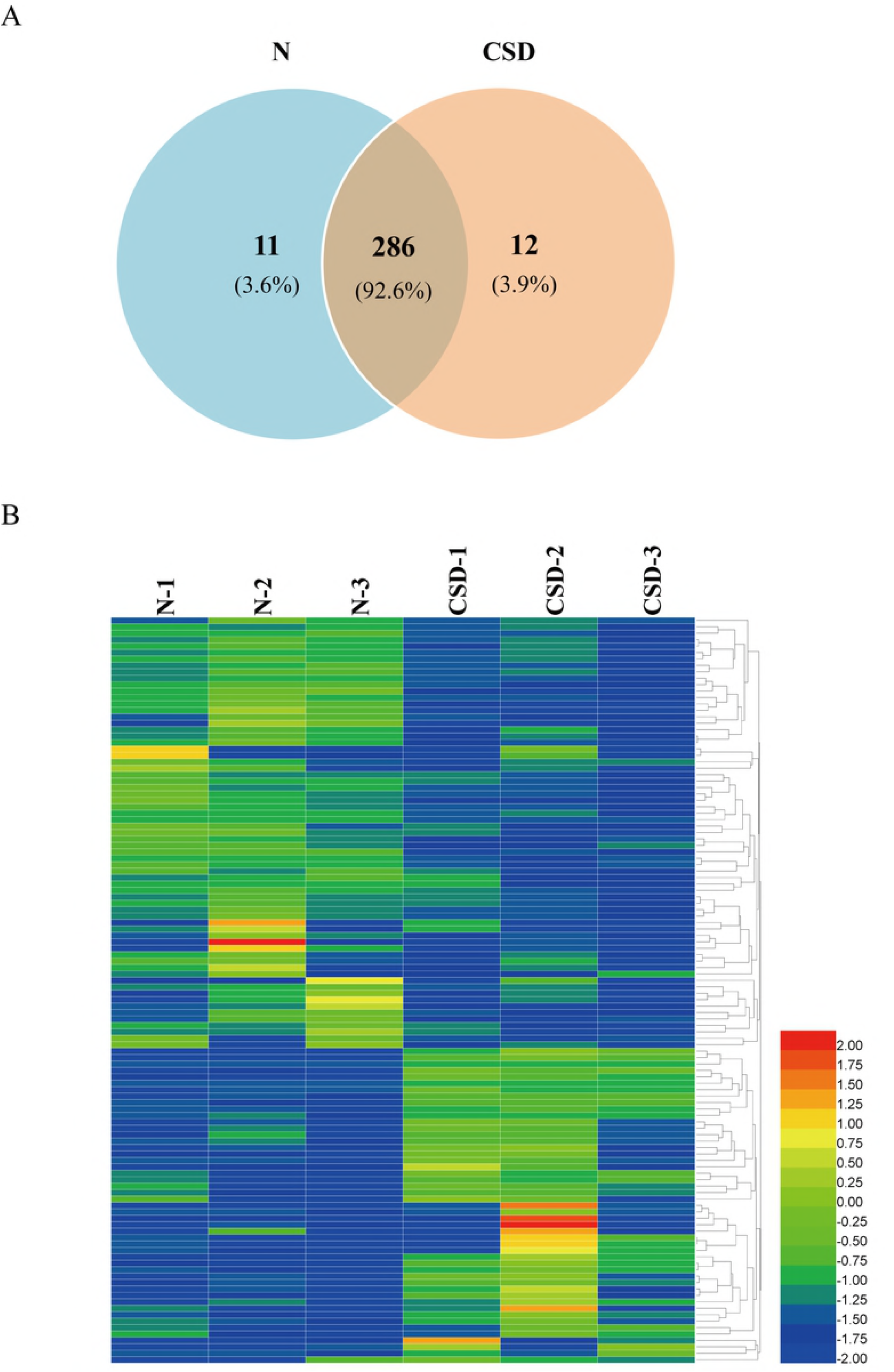
Overview of the DAPs between N- and CSD-groups. (A) Area-proportional Venn diagram depicts the overlap of the identified serum proteins between N- and CSD-group rats from mass spectrometry measurements. A total of 309 non-redundant proteins with 297 and 298 proteins were identified in N- and CSD-groups, respectively. 286 proteins were found in both groups. (B) A heat map analysis of the DAPs between N- and CSD-group. 117 proteins displayed more than 1.8-fold quantitative alteration, of which 49 proteins were upregulated and 68 were downregulated in the CSD-group compared to that of the N-group. Color depth from blue to red indicates the intensity detected by LC-MS analysis from low to high.

Label-free LC-MS/MS quantification was used to characterize the differential abundance of proteins in the serum samples. Collectively, 117 proteins displayed more than 1.8-fold quantitative alteration, of which 49 proteins were upregulated and 68 were downregulated in the CSD-group compared to that of the N-group. This analysis revealed significant changes in serum protein abundance between the two groups. A biological heat map of clusters from two groups was constructed using the raw data providing an overview of the distribution of expressed serum proteins (Figure 2B).

### Functional enrichment of the proteins with differential abundance between N-and CSD-group

To probe into the molecular mechanisms of CSD, the differential abundance proteins (DAPs) were categorized into various biological processes and molecular function classes based on Gene Oncology (GO) classification system and UniProt database. GO analysis revealed the protein classes and molecular pathways of the DAPs (89 proteins out of the 117 DAPs mapped). The protein classes include defense/immunity protein, enzyme modulator, hydrolase, signaling molecule, cytoskeletal protein, and so on (Figure 3A). And the DAPs participate in a variety of biological pathways including glycolysis, blood coagulation, integrin signaling pathway, Parkinson disease, Huntington disease, et al (Figure 3B). Additionally, functional enrichment of DAPs based on GO analyses, STRING database and UniProt database demonstrates a correlation between CSD and several physiological functions. As shown in figure 3C, the DAPs following CSD are associated with the biological processes of metabolic process, cardiovascular function, nervous system function, response to stimulus, response to stress and so on. Details of the DAPs are listed in Table s1.

**Figure 3.**
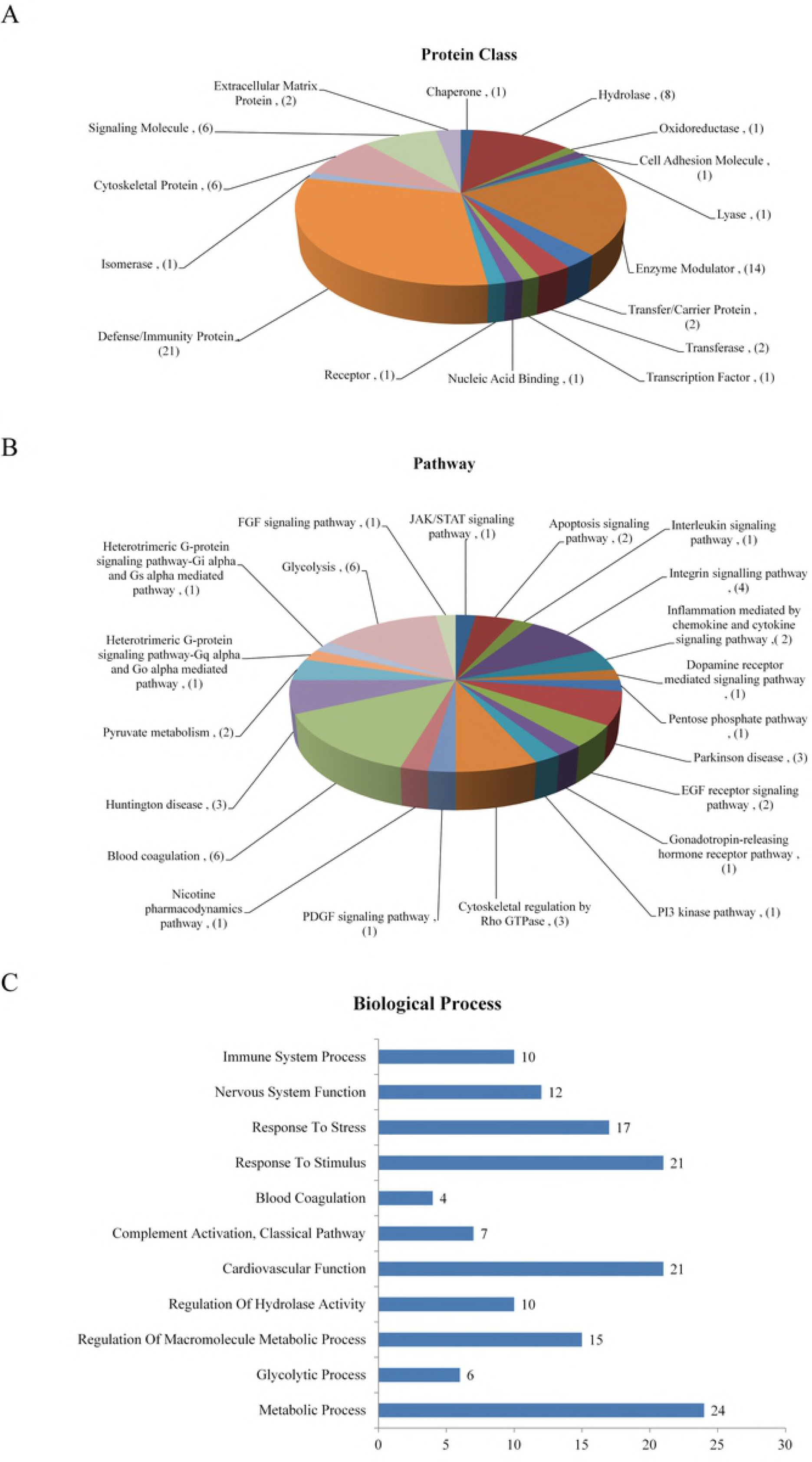
Bioinformatics analysis of the DAPs. (A & B) Functional assignments of protein class and molecular pathway of the DAPs according to gene ontology analysis (89 proteins out of the 117 DAPs were mapped). Numbers in the brackets represent the number of proteins. The protein classes include defense/immunity protein, enzyme modulator, hydrolase, signaling molecule, cytoskeletal protein, and so on. And the DAPs participate in a variety of biological pathways including glycolysis, blood coagulation, Integrin signaling pathway, Parkinson disease, Huntington disease, et al. (C) Biological and molecular function enrichment of DAPs is classified according to GO analyses and UniProt database. The bar chart shows the number of proteins in each functional class. The DAPs following CSD are associated with biological processes of metabolic process, cardiovascular function, nervous system function, and some other processes as response to stimulus, response to stress and so on.

### Protein interaction networks demonstrated two clusters involving in energy metabolism and cardiovascular function

To investigate the biological interactions among the DAPs, protein-protein functional network was constructed using STRING database (Figure 4A). 73 proteins out of the 117 DAPs were matched and several proteins showed complex interactions with other proteins. The protein functional partnership demonstrated a system-level insight into the effects of CSD. Two protein clusters were demonstrated to be highly related to CSD. The first group consists of proteins relating energy metabolism, including ENO2, GPI, CKB, PGK2, PFN1, PGAM1, PGAM2, PKLR, PKM, etc. The second group of proteins including KNG1C1S, MBL1, CLU, F12, PF4, SERPIND1, SERPING1and MMRN1, were found to be associated with cardiovascular function. The interaction network indicates that sleep plays a crucial role in maintaining the normal functions of energy metabolism and cardiovascular systems.

**Figure 4.**
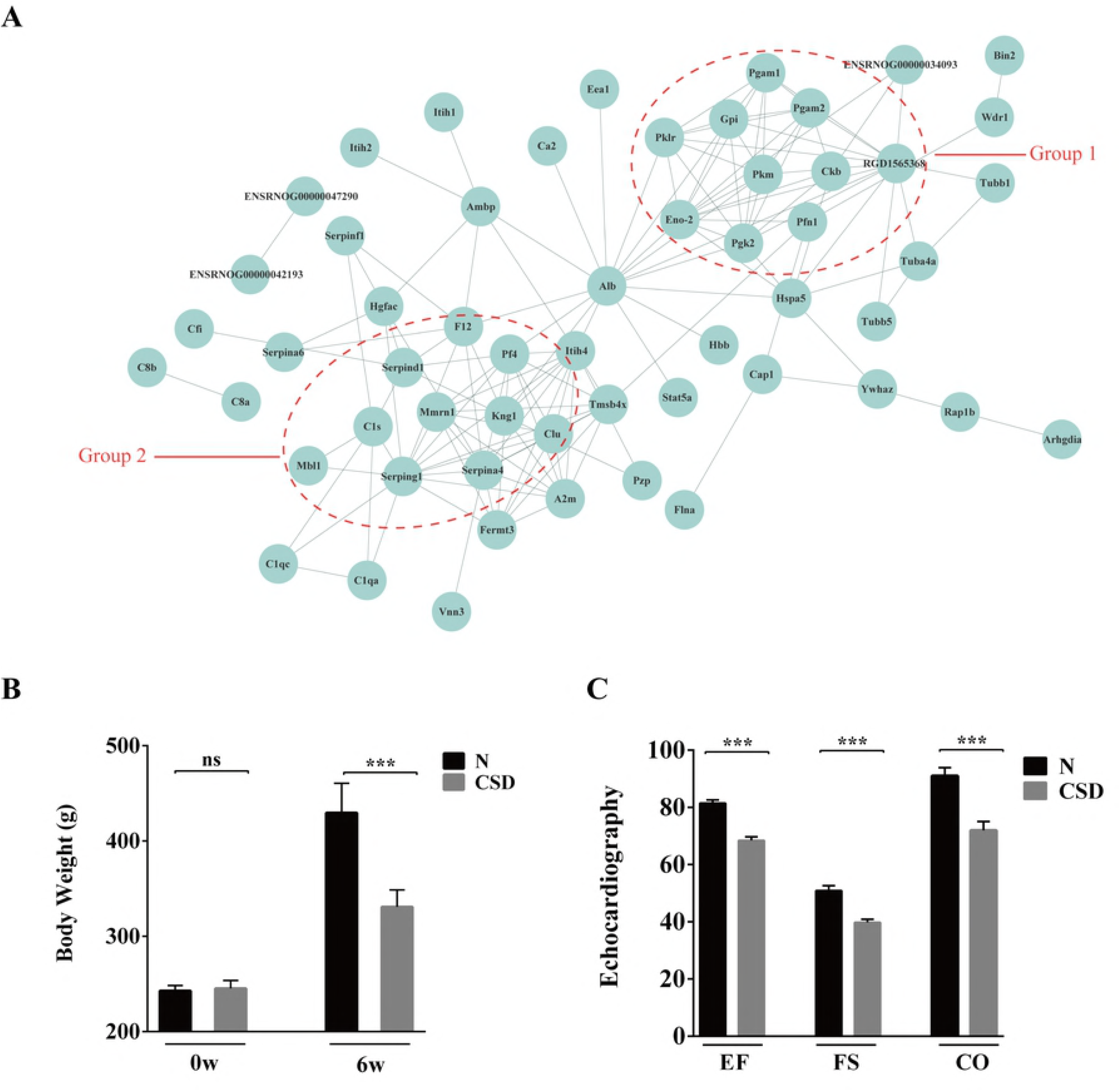
Protein interaction networks of DAPs demonstrated two clusters involving in energy metabolism and cardiovascular function. (A) STRING analysis of protein interaction networks of the DAPs. Groups 1 and 2 marked in red represent proteins involved in energy metabolism and cardiovascular function, respectively. (B) Comparison of body weight on the beginning (0w) and 6th weeks (6w) between N- and CSD-group. On the 6th week, body weight of the CSD-group rat was obviously lower than that of the N-group. (C) Echocardiography evaluation of the two groups of rats. EF: ejection fraction, FS: fraction shortening, CO: cardiac output. Myocardial function of the CSD-group was definitely weaker than that of the N-group.

Comparison of body weight on 6th weeks showed notable difference between N-and CSD-group (Figure 4B), indicating that CSD may increase energy expenditure or intensified catabolism. What’s more, echocardiography evaluation of the two groups of rats was detected using a high frequency ultrasound technology. As shown in Figure 4C, left ventricular ejection fraction (LVEF), fraction shortening (FS) and cardiac output (CO) of the N-group were significantly better than that of the CSD-group. And posterior wall of the left ventricle was significantly thickening in the CSD-group rat (Figure s1), indicating that myocardial function of the CSD-group was definitely damage compared with that of the N-group.

Collectively, the results revealed that CSD may result in abnormalities in energy metabolism and cardiovascular system.

### Identification and verification of the candidate proteins involving in CSD

Based on the bioinformatics analysis and the biological function of the differential proteins, several protein candidates were chosen to determine their expression alterations. Expression levels of the proteins were evaluated by immunoblotting assay using serum samples from external rat models. The immunoblotting assay demonstrated three proteins KNG1 (Kininogen1), PFN1 (Profilin-1), and PKM (Pyruvate kinase M1/M2) were strongly expressed in the CSD group compared to that of the control group where little or no immune recognition was found. On the other hand, one protein CLU (Clusterin) was clearly detected in the N-group, whereas its expressions were weak or undetectable in that of CSD-group (Figure 5A and B). While the remaining candidates were either undetected or showed no significant difference between the two groups.

**Figure 5.**
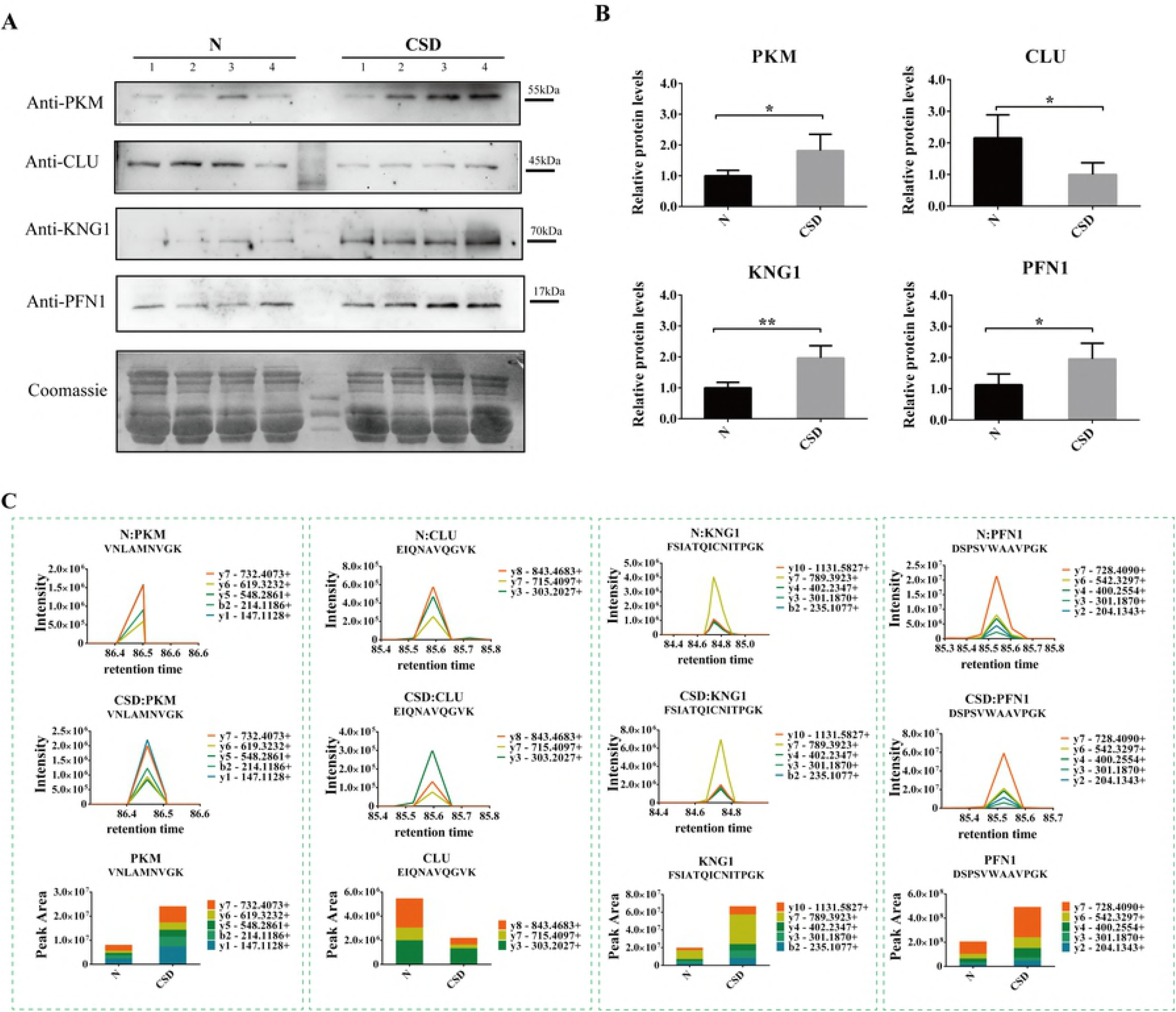
Identification and verification of the candidate proteins involving in CSD. (A) Western blots display the protein levels of four proteins PKM, CLU, KNG1 and PFN1 in N- and CSD-groups. Coomassie blue staining of the membranes were used as loading control. (Coomassie blue staining of the four proteins were shown in Figure s2.) (B) The relative intensity of the Western blot bands. KNG1, PFN1 and PKM were expressed stronger and CLU was expressed weaker in CSD-group compared with N-group. The data are reported as the mean ± SD of three independent experiments. P < 0.05 was considered significant. (C) Serum levels of the four candidates protein detected by PRM. Expression levels of PKM, KNG1 and PFN1were increased while CLU was decreased in the CSD-group, compared with the N-group.

Additionally, targeted proteomic study (PRM) was used to determine the abundance alternation of the four candidate proteins of serum samples between N- and CSD-groups. As illustrated in Figure 5C, expression levels of PKM, KNG1 and PFN1were significantly increased in response to CSD treatment. While expression level of CLU was decreased in the CSD-group.

The results confirmed a close link between CSD and alterations of the four proteins.

### Validation of the four candidate proteins in myocardial and brain tissues

Based on the above results, PFN1, KNG1, PKM and CLU may be potential candidates of CSD. As the four proteins are involved in cardiovascular and nervous system processes, myocardial and brain tissues of rat models were used for proteomic comparison using LFQ verification. As Figure 6A and B showed, expression alterations of the four proteins were highly consistent with previous results. Except for KNG1 that was not determined in brain tissue of either the two groups.

**Figure 6.**
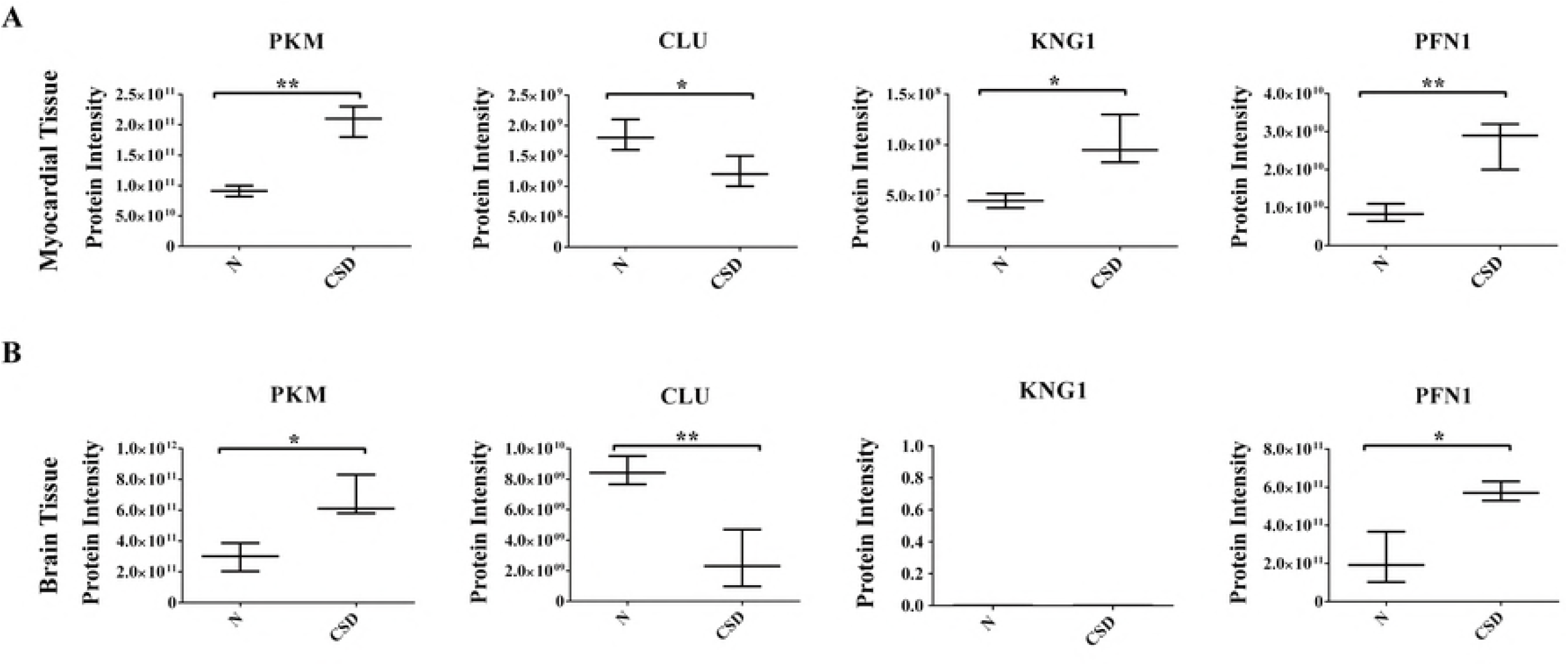
Validation of the four candidates in myocardial and brain tissues. Proteomic comparison of PKM, CLU, KNG1 and PFN1 were detected using myocardial tissue (A) and brain tissue (B) samples by LFQ verification.

Collectively, the bioinformatics analyses and validations revealed a close relationship between CSD and dysfunction of metabolic, cardiovascular and nervous systems. And comprehensive proteomics analysis revealed four proteins PKM, CLU, KNG1 and PFN1 to be potential serum biomarkers for CSD related diseases.

## Discussion

In the present study, we used a label-free quantitative proteomics technology to survey the global changes in serum protein abundance between the CSD rats and the normal rats to further elucidate the mechanisms underlying sleep disorder and relating diseases. Comparison analysis revealed a significant quantitative alteration in protein with differential abundance between the two groups. Bioinformatics and functional analyses showed a strong link between CSD and energy metabolism, cardiovascular function and nervous system. In addition, four protein candidates including PKM, CLU, KNG1 and PFN1 were identified as potential biomarkers for sleep disorder related diseases.

### Effect of CSD on energy metabolism

In this study, significant quantitative alterations in several proteins (e.g., PFN1, PKM, ENO, PGK, PGAM, GPI, PKLR, CKB) associated with energy metabolism were identified in response to CSD. PFN1 is an actin binding protein. It promotes actin polymerization by catalyzing the exchange of actin-bound ADP for ATP and transporting ATP-G-actin to the barbed end of actin^[35]^. PKM is a rate-controlling enzyme of the glycolytic pathway and catalyzes the direct transfer of phosphate from phosphoenolpyruvate to ADP to produce ATP and pyruvate^[36, 37]^. ENO (Alpha-enolase) is a glycolytic enzyme that occupies a key position in the metabolic pathway. PGK1 (Phosphoglycerate kinase 1) is an ATP-generating glycolytic enzyme in the glycolytic pathway. PGK1 catalyzes a crucial step of glycolysis, transferring a phosphate group from 1, 3-biphosphoglycerate to ADP, forming ATP and 3-phosphoglycerate^[38, 39]^. PGAM (Phosphoglycerate mutase) catalyzes the interconversion of 3-phosphoglycerate and 2-phosphoglycerate during glycolysis and plays an important role in coordinating glycolysis^[40]^. The changes in these protein levels may imply the impairments of the normal energy metabolism in response to CSD.

The current study demonstrated weight loss in the CSD animals (Figure 4B); this is consistent with that of several other animal experiments^[4]^. It has been suggested that the CSD was linked to food intake, body weight and energy expenditure, metabolic syndrome and glucose homeostasis^[4]^. Additionally, CSD rat models showed several metabolic alterations such as increased energy expenditure and intense catabolism.

### Effect of CSD on cardiovascular system

Sleep is a vital regulator of cardiovascular function, both in the physiological state and in disease conditions^[41]^. Sleep disorder exerts harmful effects on a variety of body systems due to pathological factors such as arrhythmia, sympathetic activation, high blood pressure, oxidative stress and endothelial dysfunction^[42]^. In this study, several proteins associated with cardiovascular function (such as KNG1, C1S, PFN1, PKM, ENO1, MMRN1, CLU, SERPIND1, PF4, F12 and FLNA) showed abundance alterations. KNG1 is an inflammation mediator (KNG1) that inhibits the thrombin- and plasmin-induced aggregation of thrombocytes^[43]^. As an actin binding protein, PFN1 is also involved in the atherosclerotic vascular and cardiac hypertrophy diseases^[44, 45]^. PKM also takes part in cardiac function through participation in energy metabolism regulation of the cardiac muscle cells^[37, 46, 47]^. ENO1 plays a catalytically independent role in cardiomyocyte apoptosis. It participates in the protection of the cardiomyocytes against oxidative stress and play roles in the pathogenesis of cardiac hypertrophy^[48, 49]^. MMRN1 (Multimerin 1) is a homopolymeric protein that is stored in platelets and endothelial cells for activation-induced release. It supports the adhesion of many different cell types including activated platelets, neutrophils, and endothelial cells^[50]^. As a normal constituent of the vascular sub endothelial matrix, MMRN1is proposed to support platelet function at sites of vessel injury. It supports platelet adhesive functions and thrombus formation^[51]^. CLU is a secreted chaperone protein. Clinical research has suggested that increasing of CLU level in blood during cardiovascular disease might limit the damage and lead to a more favorable prognosis for patients through saving reversibly damaged cardiomyocytes^[52, 53]^. PF4 (Platelet factor 4) is a very abundant platelet α-granule CXC chemokine that is released during platelet activation^[54, 55]^. Several investigations have reported that PF4 exerts a number of effects on various aspects of hemostasis and thrombosis. But the effects are pleiotropic because of its high affinity for negatively charged molecules rather than binding to a single or small number of specific receptors^[54]^. F12 (Factor XII) is a procoagulant factor. It plays an essential role in blood coagulation and has a profound influence on thrombin generation^[56, 57]^. Given the key role of F5 in hemostasis, it was considered as a risk factor for thrombosis and hemophilic clotting disorders^[58]^. FLNA (Filamin-A) was encoded by a familial cardiac valvular dystrophy gene. It plays crucial role during vascular development and cardiac morphogenesis^[59, 60]^. Mutations in FLNA may result in cardiac valvular dystrophy, aneurysms, cardiac defects and neurological dysfunction^[61, 62]^.

The important roles of sleep in the cardiovascular system have been highlighted in recent years. Clinical research revealed that CSD is associated with the morbidity and mortality of coronary heart disease. Echocardiography evaluation of the CSD group rats in the current study also showed significant abnormality (Figure 4C), suggesting that CSD may exert harmful effects on cardiovascular system.

### Effect of CSD on nervous system

CSD affects a wide network of brain structures and is a significant causative factor in the development of neurodegenerative diseases such as Alzheimer’s disease^[63]^. ENO1 is a multifunctional protein that plays a role in Alzheimer’s pathology^[52, 53]^. CLU has been shown to be clearly upregulated in astrocytes of the Alzheimer’s disease patients^[53, 64]^. KNG1, an inflammation mediator may be involved in neuronal damage leading to neuronal diseases^[65]^. Our data of serum protein alterations indirectly indicate the potential effects of CSD on nervous system function which may be associated with several neurodegenerative diseases.

In summary, the alterations of plasma protein abundance induced by CSD may have clinical and biological relevance with the dysfunction of metabolic, cardiovascular and neurological systems.

### Molecular and biological functions of the four potential biomarkers of CSD

In the current study we identified four proteins including KNG1, PFN1, PKM and CLU that are highly associated with CSD. These proteins may serve as potential serum biomarkers for further pathological and clinical research of the disease.

The inflammation mediator KNG1 belongs to the plasma kallikrein-kinin system, which is an important regulatory system in body defense mechanisms such as inflammation and blood pressure control^[43]^. KNG1 is a central constituent of the contact-kinin which represents an interface between thrombotic and inflammatory circuits. It inhibits the thrombin- and plasmin- induced aggregation of thrombocytes. Bradykinin, the active peptide that is the cleave product of KNG1, shows a variety of physiological effects such as induction of hypotension, decreased blood glucose level and increased vascular permeability^[43, 65–67]^. Additionally, investigations based on KNG1 knock out mice suggested that KNG1 may be associated with neuronal damage via different pathways involving in activation of the contact-kinin system: enhanced microvascular thrombosis, blood-brain barrier leakage, and inflammation^[65]^. In our study, KNG1 was upregulated in the CSD-group. Higher expression of KNG1 induced by CSD may result in higher-risk of cardiocerebral vascular and neuronal system diseases.

PFN1 is an actin binding protein. Traditionally, PFN1 has been considered as a key regulatory protein for actin polymerization and cell migration contributing to many biological activities through assembling and disassembling actin filaments^[68]^. Results from *in vivo* experiments revealed that PFN1 plays an important role in chondrodysplasia, cerebellar hypoplasia and endothelial proliferation^[69–71]^. In addition, loss expression of PFN1 on mammalian cell systems may cause impaired migration/invasion and capillary morphogenesis of human vascular endothelial cells, and defects in neurite outgrowth^[72,73]^. Several studies have suggested PFN1 to be a critical promoter of cardiac hypertrophy. Expression of PFN1 could inhibit the expression of caveolin-3 and the activity of eNOS/NO pathway, exerting a promotion effect to the development of hypertensive cardiac hypertrophy^[74]^. PFN1 expression is significantly enhanced in human atherosclerotic plaques and the serum levels of PFN1 correlate with the degree of atherosclerosis in humans^[44, 45]^. PFN1 is secreted from fully activated platelets in the thrombotic mass and it can be detected in the systemic circulation, making it a potential marker of ongoing thrombosis and of the elapsed time of ischemia^[75]^. Overall, these studies revealed complex functions of PFN1 as a modulator of sarcomeric organization and as a mediator of hypertrophic cardiomyocyte remodeling. In this study, PFN1 was upregulated in the CSD group, suggesting elevated level of PFN1 in response to CSD may be related to the cardiac dysfunction of the rat model. Therefore, PFN1 may serve as a potential useful biomarker for sleep disorders and may also represent an important therapeutic target for the treatment or prevention of the CSD-related cardiovascular diseases.

PKM is a rate-controlling enzyme of the glycolytic pathway modulating the final step of glycolysis and intracellular signaling inputs with the metabolic state of the cell^[76]^. Previous investigations revealed the significance of PKM in metabolism process, cell proliferation and cardiac function^[47, 77]^. Pyruvate kinase consists of four isoforms in mammals. Among them, PKM2 is the only one to be allosterically regulated between an active tetramer and a less active dimmer^[78]^. The ability of PKM2 to rapidly cycle between a tetramer and dimer could be especially advantageous to the failing heart^[46]^. A strong coupling between endogenous pyruvate kinase and ATPase was demonstrated in cardiac muscle cells highlighting the importance of glycolysis in energy production for cardiac function^[36, 37]^. Examination of tissue from patients with heart failure displayed a shift in the enzyme pyruvate kinase from the PKM1 isoform to the PKM2 isoform. This may be consistent with that a salient feature of the failing heart is metabolic remodeling towards predominant glucose metabolism^[46]^. In our study, the increased expression of PKM in response to CSD indicates that a higher level of energy consumption may be needed in the sleep-deprived rats to maintain their activities. And PKM may represent an important novel target for metabolic and cardiovascular diseases associated with sleep disorders.

CLU is a secreted chaperone protein expressed by a wide array of tissues. It is involved in various versatile physiological processes including cell adhesion, spermatogenesis, cell-cycle regulation, tumor metastasis, etc. The expression level of CLU may be induced by diverse conditions of cell stress and tissue injury, including myocardial infarction, ischemia, inflammation, apoptosis, and oxidative stress^[79]^. Experiments in mouse model of myocarditis revealed that CLU expression was dramatically upregulated in ventricular myocytes, especially in bordering areas of inflammation and myofiber atrophy. It has also been suggested that CLU limited progression of autoimmune myocarditis and protected the heart from post inflammatory tissue destruction^[80]^. Decreased levels of plasma CLU were associated with progression of chronic heart failure, suggesting that CLU have cytoprotective properties and might be capable to prevent myocardial injury^[81]^. Furthermore, it has been suggested that CLU exerts the neuroprotective effects *via* preventing excessive inflammation, inhibiting the activation of the complement system and cleaning of dead material^[64]^. Injection of CLU to the brains of Alzheimer’s disease mice has demonstrated anti-inflammatory and anti-apoptotic effects^[52]^. In this study, CLU was downregulated in the CSD-group, suggesting that CSD may affect the normal functions of cardiovascular and nervous system.

### Conclusion

The results of the present study demonstrate significant alterations of serum protein abundance in response to CSD. Functional and bioinformatics analyses revealed a close link between CSD and several biological processes including energy metabolism, cardiovascular function and nervous function. Expression of four proteins including PKM, CLU, KNG1 and PFN1 were changed by CSD and these proteins can potentially serve as biomarkers for sleep disorders. However, because energetics/metabolism dysfunction may affect the normal function of many other cells in the body, further studies on cardiac/brain tissue and clinical samples are necessary to confirm whether the proteome changes are specific to CSD. Nevertheless, the current study provides insights into the pathological and molecular mechanisms underlying sleep disorder at protein level.

---

**Figure s1**. Transthoracic echocardiography evaluation of two groups of rats. Posterior wall of the left ventricle was significantly thickening in the CSD-group rat.

**Table s1**. Details of the DAPs of rat serum LC-MS analysis.

**Table s2**. Peptide sequences of the four candidates verified in PRM study.

**Supplementary data 1**. Details of the LC-MS data of rat serum.

**Supplementary data 2**. Details of the LC-MS data of rat cerebral tissue.

**Supplementary data 3**. Details of the LC-MS data of rat myocardium.

## Declarations

### Ethical approval

The study was approved by the Ethics Committee of Xi Yuan Hospital, China Academy of Chinese Medical Sciences.

### Availability of data and material

The datasets used and/or analyzed during the current study are available from the corresponding author on reasonable request.

### Authors’ contributions

J-C. C. developed the animal models; J-C. C. and M. B. designed the experiments; M. B., Y-Y. M. and D-P. W. performed LC-MS and bioinformatics analysis; M. B., B-J. X. and A-M. Z prepared figures; M. B.,Y-H. P, D.C. and J-X. L. wrote the manuscript. All authors read and approved the final manuscript.

### Funding

The current thesis was supported by National Natural Science Foundation of China (81573650), the National Key Basic Research Special Foundation of China (2015CB554405) and the Chinese Postdoctoral Science Foundation (2017M621041).

### Conflicts of interest

The authors declare that they have no conflicts of interest.

### Consent for publication

This manuscript is solely submitted to Clinical Proteomics for consideration.

